# Re-evaluating the conventional wisdom about binding assays

**DOI:** 10.1101/2020.02.03.932392

**Authors:** Brandon D. Wilson, H. Tom Soh

**Affiliations:** Department of Chemical Engineering, Stanford University, Stanford, CA 94305, USA; Department of Electrical Engineering, Stanford University, Stanford, CA 94305, USA; Department of Radiology, Stanford University, Stanford, CA 94305, USA; Chan Zuckerberg Biohub, San Francisco, CA 94158, USA

**Keywords:** Binding assays, Affinity reagent, Digital detection, Langmuir isotherm, Specificity

## Abstract

Analytical technologies based on binding assays have evolved substantially since their inception nearly 60 years ago, but our conceptual understanding of molecular recognition has not kept pace. Indeed, contemporary technologies such as single-molecule and digital measurements have challenged, or even rendered obsolete, core aspects of the conventional wisdom related to binding assay design. Here, we explore the fundamental principles underlying molecular recognition systems, which we consider in terms of signals generated through concentration-dependent shifts in equilibrium. We challenge certain orthodoxies related to binding-based detection assays, including the primary importance of a low *K*_D_ and the extent to which this parameter constrains dynamic range and limit of detection. Lastly, we identify key principles for designing binding assays optimally suited for a given detection application.

## Molecular Quantification via Concentration Dependent Shifts in Equilibrium

The first published implementation of a binding-based molecular assay was an **immunoassay** for insulin detection in 1960 [1]. Over the following 60 years, analytical technologies based on **binding assays** have evolved substantially. Molecular detection platforms based on enzyme-linked immunosorbent assays (ELISAs) as well as newer commercial technologies, such as Luminex and NanoString, are core components of today’s diagnostic armamentarium. In general, binding assays generate signal through concentration-dependent shifts in equilibrium that result from the interaction of the target molecule with an **affinity reagent**. Typically, either the target or affinity reagent is directly coupled to a moiety that generates an observable readout upon binding. For example, myriad immunosensors [2] have been developed that couple the interaction between antibody and antigen to an observable output based on an electronic [3], optical [4], or mechanical [5] signal, or even on the production and amplification of a quantifiable DNA sequence [6]–[8]. In all of these cases, an increase in target concentration shifts the equilibrium toward the formation of more antibody-antigen complexes, increasing the signal. Microarray technology likewise quantifies gene expression in a manner where the equilibrium fraction of target molecules bound to the array surface—and thus the fluorescence signal generated at their cognate array feature—is proportional to their concentration in the bulk solution [9]–[11].

A conventional understanding of molecular recognition principles has been used for decades to describe microarray and immunoassay technologies. However, the advent of contemporary molecular detection technologies, such as single-molecule measurements [12]–[16], has rendered important aspects of this understanding either unnecessarily restrictive or even obsolete. For instance, it is commonly assumed that the useful **detection range** of an affinity reagent spans an 81-fold change in target concentration centered around the equilibrium **dissociation constant** (*K*_D_). This heuristic is still useful for conventional immunoassays, but it may be overly limiting in the broader context of molecular detection assays because it implies that one can only quantify target concentrations as low as *K*_D_/9. There is also a prevalent notion that a lower *K*_D_ is always “better” because it results in a lower **limit of detection** (LOD), however there are many scenarios in which this would be detrimental to the assay. This is not to say that the conventional wisdom about molecular recognition is wrong, but rather that stringent adherence to prior conventions limits what could otherwise be achieved with newly available technologies. Here, we aim to reevaluate the conceptual frameworks used when developing binding assays and to provide an intuitive understanding of molecular recognition, empowering researchers to make better use of contemporary technologies for quantitative analysis of biological systems. A number of excellent articles have already shown how to shift and manipulate binding curves from a physicochemical perspective [17]. Therefore, we endeavor to approach the topic from a stance that is agnostic to the specific assay chemistry or implementation. After providing a brief background on the fundamental principles of affinity-based assays, we use simple mathematical reasoning and literature examples to argue against some long-held preconceptions of molecular recognition.

## Single-site molecular recognition

Although more complicated examples of molecular recognition such as population shift [18]–[20], allostery [21]–[23], and proximity assays [24] have proven extremely powerful, many novel insights about assay development can still be gleaned from a simple model of bimolecular association. In the standard model of bimolecular association, the binding strength between an affinity reagent, *A*, and a target molecule, *T*, is characterized by the binding affinity, which is quantified in terms of *K*_D_ (eq. 1).

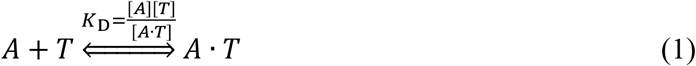

Binding affinity plays a crucial role in determining what range of target concentrations yields an observable signal (**Figure 1A**). A typical antibody-antigen interaction has a *K*_D_ of ~1 nM [25]. An affinity reagent is generally considered high affinity when *K*_D_ < 10 nM and low affinity when *K*_D_ > 1 μM, although, as we will discuss, this is not necessarily the best conceptual approach. The **binding curve** is commonly defined in terms of the Langmuir isotherm [26]–[28], an equation that characterizes the amount of binding as a function of target concentration (eq. 2). It is not without limitation (Box 1.). Several key assumptions underlie the simplicity of the Langmuir isotherm: i) single-site binding, ii) single *K*_D_, iii) one component is in excess, and iv) no off-target reactions. If these assumptions hold, one can derive the fraction of affinity reagent bound to target as

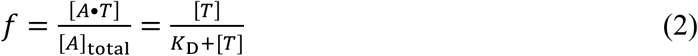

for conditions in which [A] ≪ [T]. This simple equation has informed much of our conventional understanding of molecular recognition, e.g. that 50% of affinity reagent molecules are bound to target when the target concentration is equal to the *K*_D_ (**Figure 1B**), which defines the center of the dynamic range around the affinity. In this context, the useful detection range has conventionally been defined as the range of target concentrations at which 10–90% of the affinity reagents are bound to target [17]. A cocktail-napkin analysis reveals that the detection range in this model spans target concentrations from 0.11–9.0 x *K*_D_, representing an 81-fold concentration window (**Figure 1b**). This implies that an affinity reagent with a *K*_D_ of 1 nM can only quantitatively detect targets in the range of 111 pM to 9 nM. However, the utility of this heuristic is called into question by the myriad examples of affinity reagents being used to quantify target concentrations far below their *K*_D_.

**Figure 1.**
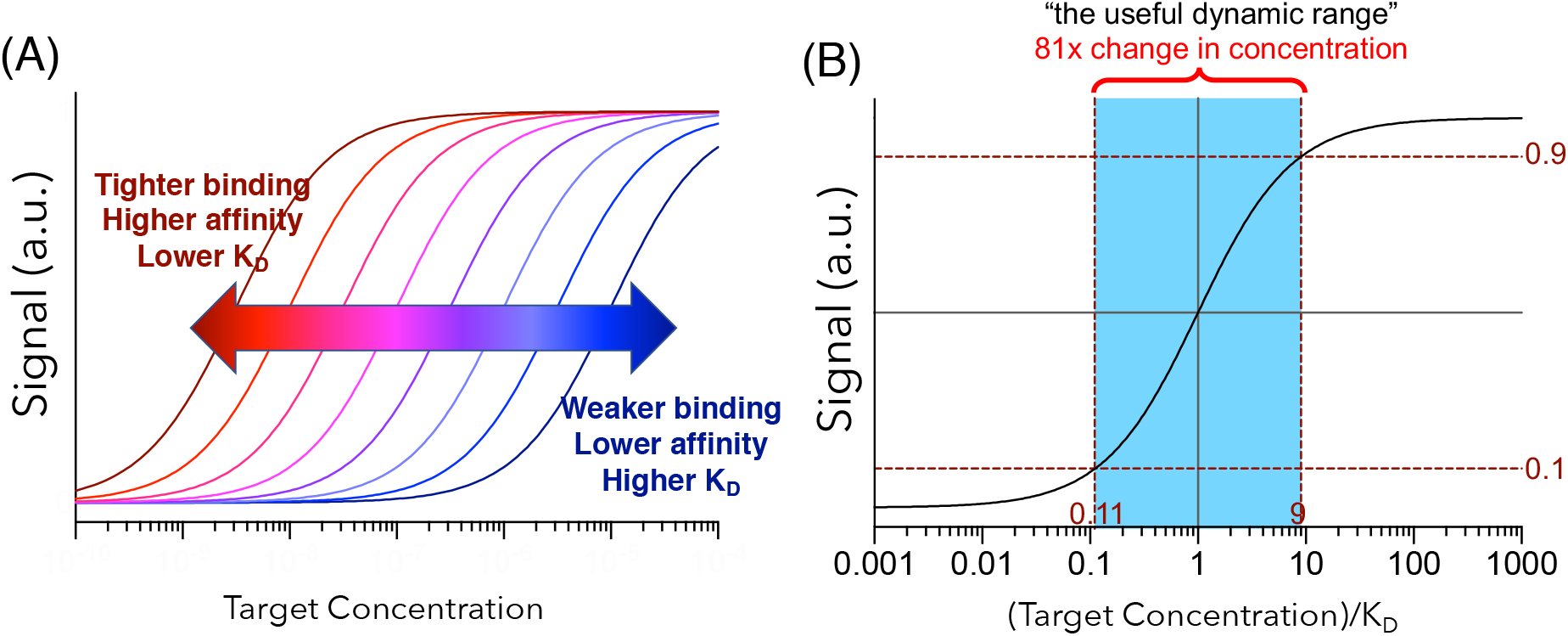
Conventional understanding of binding curves. (**a**) Increasing affinity (*i.e.*, lowering *K*_D_) shifts binding curves leftward relative to target concentration, whereas lowering affinity (*i.e.*, increasing *K*_D_) leads to a rightward shift in the binding curve. (**b**) The “useful detection range” (blue) is considered to encompass the target concentrations over which 10–90% of the affinity reagent molecules are target-bound. This yields detectable target concentrations that span 0.11– 9.0 x *K*_D_.

#### Box 1. When can we use the Langmuir isotherm and when does it fall apart?

A core – but often overlooked – assumption enabling the use of the Langmuir isotherm is that one component of the system must be in vast excess of the other so binding does not cause depletion of one of the species. If the target is much more abundant than the affinity reagent (*i.e.,* [*T*] ≫ [*A*]), then changes in the target concentration lead to changes in the fraction of affinity reagent that is bound to the target:

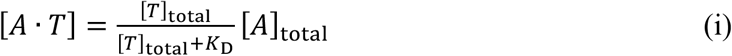

In this case, the resultant signal is non-linear in terms of target concentration but linear in terms of affinity reagent concentration. In contrast, if the affinity reagent is present at a much higher concentration than the target (*i.e.,* [*A*] ≫ [*T*]), then the fraction of bound target remains constant while the total amount of target varies:

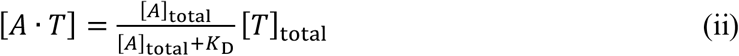

Here, the signal is non-linear in terms of affinity reagent concentration and linear in terms of target concentration. Since it is typically easier to couple a detection modality (*e.g.*, a fluorescent or bioluminescent tag) to an affinity reagent than to the target, the background signal scales with [*A*] in most experimental setups. Therefore, the affinity reagent-limited scenario usually exhibits a more favorable signal-to-noise ratio than the target-limited scenario. However, a practical consideration of imposing the condition that [*T*] ≫ [*A*] is that in order to measure low target concentrations, affinity reagent concentration must be even lower. Thus, in the absence of signal amplification, the detection limit is constrained by the detection modality rather than the affinity reagent.

The conventional Langmuir isotherm no longer holds if affinity reagent and target concentrations are comparable. In this case, the amount of target-affinity reagent complexes is a non-linear function of the concentrations of both components[49]:

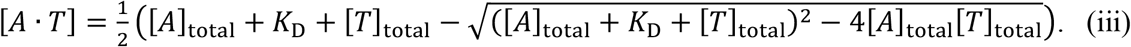

If [*A*] = [*T*] = *K*_D_, rather than observing the intuitive 50% binding, one would instead observe ~38.2% binding. The concentrations must be raised to [*A*] = [*T*] = 2*K*_D_ to achieve 50% binding.

The Langmuir isotherm can be a powerful tool for rationalizing and analyzing binding data, especially because it conveniently distills a binding curve into a single parameter. However, the misuse of the equation can lead to misinterpretations and erroneous conclusions.

## What is a “good” K_D_?

Other misconceptions related to molecular recognition stem from our own implicit biases. One such bias is the concept of an objectively “better” *K*_D_; we are biased towards thinking that a lower *K*_D_ is intrinsically superior because we predominantly work under conditions where the *K*_D_ is higher than the concentration of the target we are trying to detect. In practice, the concept of a “better” *K*_D_ is only relevant in the context of a desired detection range. Many researchers, however, fall into the trap of optimizing for lower and lower LODs—and by proxy, lower and lower *K*_D_— without regard for the biological scenario in which the assay is to be used.

We often overlook the fact that quantifying albumin concentrations in the 1 mM range using an affinity reagent with *K*_D_ = 1 μM is as challenging as – or, as we will discuss later, possibly even more challenging than – quantifying insulin concentrations in the 1 nM range using a reagent with the same affinity (**Figure 2A**). If the *K*_D_ is too high, there will be too little signal over the desired detection range; if the *K*_D_ is too low, the signal will be saturated. In either case, one loses the ability to differentiate changes in target concentration via shifts in equilibrium. What matters is matching the detection range to the clinical scenario at hand. Therefore, a good effective *K*_D_ or dynamic range can only be defined in terms of a given biological context. For example, an affinity reagent with a *K*_D_ of 1 μM is perfect for detecting ATP in blood, since its basal level is on the order of 20 nm to 1 μM[29]. In contrast, an affinity reagent with *K*_D_ = 1 pM towards ATP would be functionally useless because any reasonable deviation in ATP concentration would not result in an observable change in binding. Unfortunately, it remains common in the realm of biosensor development to report *K*_D_ without *any* mention of the clinically relevant detection range.

**Figure 2.**
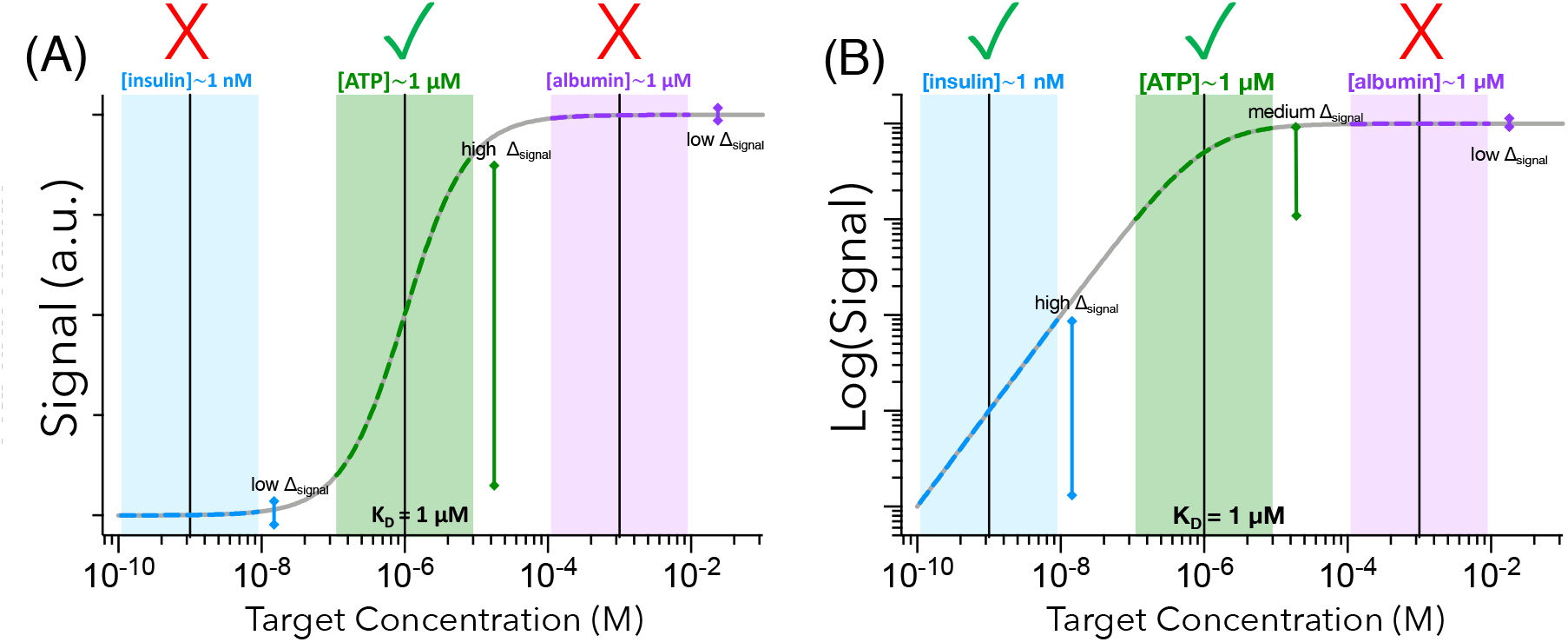
The right *K*_D_ for the job. **a)** Detecting 1 nM insulin with an affinity reagent with *K*_D_ = 1 μM is just as challenging as detecting 1 mM albumin. If the affinity is too high or too low, the output signal will not change appreciably with changes in target concentration. **b)** The greater sensitivity of contemporary single-molecule and digital detection assays means that one can, with sufficiently low background, quantitatively detect 1 nM insulin more easily than 1 mM albumin or even 1 μM ATP using an affinity reagent with a *K*_D_ of 1 μM. In this scenario, a logarithmic scale for detection is more useful than the linear scale employed in **a**.

## Logarithmic Detection Modalities and the Influence of Background Signal

The advent of sophisticated detection modalities with extremely high resolution and considerably lower levels of background should encourage a reconsideration of how best to define signal output. In contrast to traditional readouts like bulk fluorescence or electrochemistry, which report ensemble averages of the number of bound affinity reagent molecules, single-molecule and digital assays report individual molecular counts. Since these outputs are quantitative over orders of magnitude, it is more useful to define the y-axis on a logarithmic scale. This has the important result that the quantification of analytes at concentrations arbitrarily lower than *K*_D_ becomes possible, assuming zero background signal. This is in stark contrast to the conventional thinking that the functional detection range is strictly proscribed relative to the *K*_D_. Moreover, the error associated with the calculated concentration decreases with the logarithm of the signal—that is, measurements get *more* accurate at lower concentrations (**Figure 2B**). Purely digital measurements with zero background are limited solely by Poisson statistics.

In the absence of background signal, the log dynamic range extends indefinitely, retaining quantitative resolution over arbitrarily low concentrations (**Figure 3A**). This means that in theory, the ability to detect low concentrations can be completely decoupled from *K*_D_. In practice, background signal strongly influences the LOD [30] and the detection range of an assay. Background signal can often be decreased through assay design or optimization whereas binding affinity is generally—with some exceptions[17], [18], [20], [23]—a fixed property of an affinity reagent-target pair. Moreover, a background signal of 1% has a large impact on a log detection range but a negligible effect on the conventional linear range (**Figure 3B**). Therefore, the minimization of background signal is more attractive for improving LOD than relying on decreasing *K*_D_. Specifically, assuming a constant coefficient of variation (CV) for background signal, decreasing the background signal will have the same effect on the LOD as reducing *K*_D_ by the same factor (**Figure 3C**).

**Figure 3.**
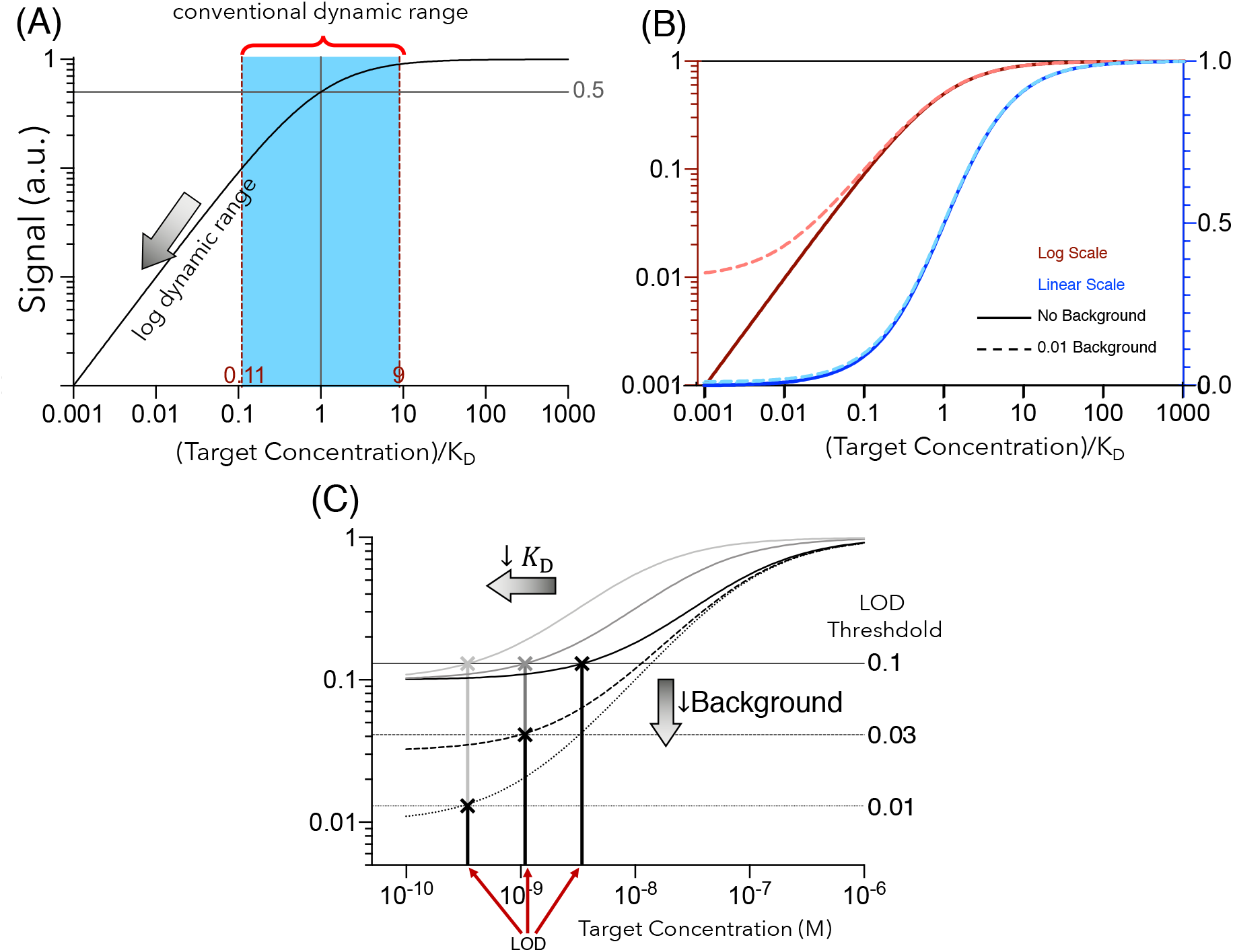
Logarithmic detection and background signal. (**A**) If the y-axis can be plotted on a log-scale, which is appropriate for detection modalities that can distinguish between small changes in binding fraction, then the log-linear detection range extends below the conventional detection range (blue) as defined in **Figure 1B**. In the absence of background signal, the log detection range includes indefinitely low target concentrations and the center of the log dynamic range is no longer located at [Target] = *K*_D_. (**B**) Background signal has a larger impact on a logarithmic output than a linear output. The extent to which the log detection range extends below *K*_D_ is a strong function of background signal. Adding a background signal of 1% does not appreciably affect the dynamic range for a linear output, but it greatly reduces the dynamic range for a log output. (**C**) Decreasing *K*_D_ (dark to light solid lines) has the same effect on LOD as decreasing the background signal by the same factor (solid to dashed to dotted black lines). Horizontal lines represent the background signals equivalent to 1, 3, and 10% plus three standard deviations, assuming 10% coefficient of variation. In contrast, *K*_D_and background signal can have different effects on the detection range. Decreasing *K*_D_ does not appreciable change the extent of the detection range, whereas decreasing background signal greatly extends the log detection range.

Sophisticated detection modalities with considerably lower levels of background and digital outputs enable us to quantify molecules at concentrations vastly below *K*_D_, thereby rendering the traditional notion of an 81-fold detection range centered around the *K*_D_ obsolete. Instead, it is more useful to define a log-linear detection range where, given a sufficiently sensitive detection modality and low enough background, it should be possible to detect analytes at concentrations arbitrarily lower than the *K*_D_. A corollary of this rationale is that the burden of optimizing LOD should not be placed solely on decreasing the *K*_D_ of the affinity reagent itself but also on the development of low-background assays with detection modalities that can discriminate small changes (*e.g.,* 0.1–0.01%) in the fraction of bound affinity reagents. Detection modalities that are capable of this level of resolution typically involve the counting of single-molecule events, such as droplet digital PCR [31], DNA sequencing [32], [33], digital ELISA [13], or single-molecule imaging [15]. With a sufficiently low background signal and a logarithmic detection modality, it is possible to detect and quantify target concentrations many orders of magnitude below *K*_D_. An interesting ramification of this rationale is that the *K*_D_ of the affinity reagent must be higher than the physiological range of target concentration in order to ensure an appropriate log-linear detection range. Indeed, it is actually *worse* to have a *K*_D_ lower than the relevant concentration range.

## A Case Study

This concept of extending the detection range through the selection of an appropriate detection modality and the elimination of background signal is exemplified by recent work from Li and colleagues [34], where they were able to consistently quantify analytes at concentrations that are orders of magnitude below *K*_D_. The authors used a three-state population-shift mechanism, which they termed recognition-enhanced metastably-shielded **aptamer** probes (RMSApt). The affinity reagents are designed in such a way that they can initiate a rolling circle amplification (RCA) reaction [35], [36] when bound to the target molecule. They are coupled to a solid surface, so that individual RCA products can be imaged and counted. This assay inherently exhibited high background signal, because aptamers not bound to target could also initiate an RCA reaction. The authors solved this problem by introducing an enzymatic “locking” mechanism that results in the non-equilibrium depletion of unbound aptamers, functionally eliminating background signal. Impressively, the authors observed quantitative detection ranges and LODs that consistently extend many orders of magnitude below the reported aptamer *K*_D_. These improvements in quantitative **sensitivity** appear to be very robust and were demonstrated with three different affinity reagents. An ochratoxin A (OTA) aptamer with a reported *K*_D_ of 500 nM [37] exhibited a LOD of 38.8 fM and a detection range of 100 fM to 1 nM (**Figure 4A**); a kanamycin aptamer with a reported *K*_D_ of ~80 nM [38] exhibited a LOD of 8.9 fM and a detection range of 10 fM to 10 pM (**Figure 4B**); and a tyrosinamide aptamer with a reported *K*_D_ of 45 μM [39] exhibited a LOD of 47.5 pM and a detection range of 10 pM to 100 nM (**Figure 4C**). The authors consistently demonstrate the quantification of concentrations orders of magnitude below *K*_D_!

**Figure 4.**
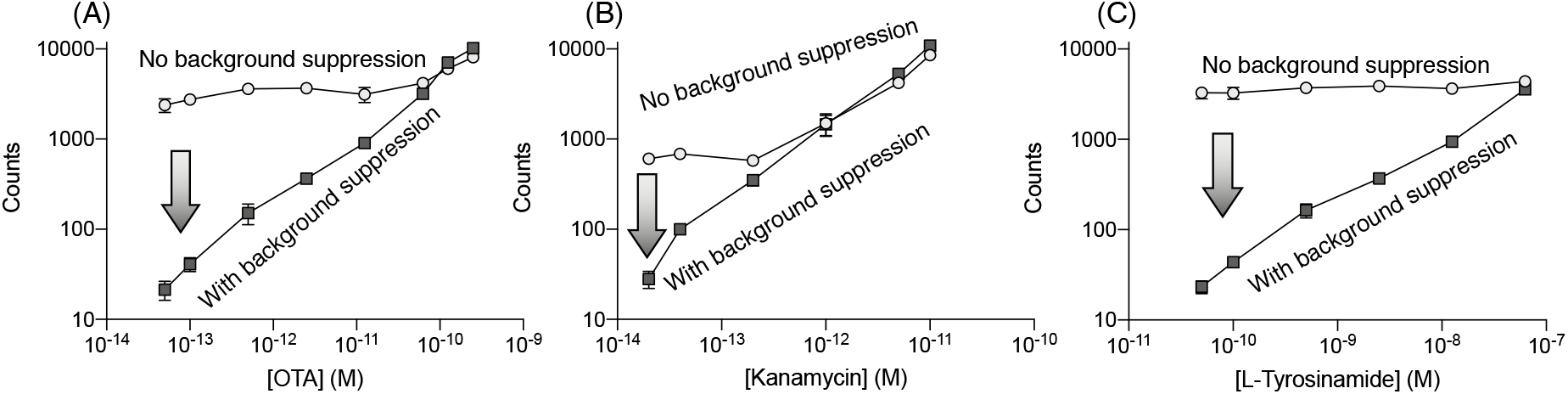
Digital detection and low background single render the concept of an 81-fold detection range centered around the *K*_D_ obsolete. The recognition-enhanced metastably-shielded aptamer probe (RMSApt) system [34] exemplifies how a digital readout with little to no background signal on a log output scale can dramatically extend an assay’s functional detection range. RMSApt essentially eliminates background signal through an enzymatic locking mechanism, achieving LODs and quantitative detection ranges that are orders of magnitude below reported affinities of the aptamers for their targets: **a**) ochratoxin A (OTA), **b**) kanamycin, and **c**) L-tyrosinamide. Importantly, a digital readout by itself (circles) is not sufficient to achieve these results; rather, a mechanism of background suppression (squares) must also be included. Data has been replotted from Ref. 36 with permission from the authors.

Although these results initially seem impossible in light of traditional understanding of molecular recognition, they should not be surprising considering the impact that background signal and the use of a digital detection modality have on the functional detection range. The high resolution is conferred in part by the ability to image and count individual binding events, which enables discrimination between small changes in the fraction of bound aptamers, and in part by the functional elimination of background through an enzymatic locking mechanism, which enables detection at concentrations far below *K*_D_. In this scenario, the ability to distinguish between small percentages of binding events is limited only by the number of viewable RCA products—which is in turn constrained by Poisson noise at the low end of target concentration and by image saturation at the high end. Thus, the ability to perform a high-sensitivity assay should not be exclusively dictated by access to high-affinity binding reagents but should also exploit a high-resolution detection modality and the ability to minimize background signal.

The previous example required the introduction of an irreversible reaction to the assay. This is not a necessity; many other methods can achieve similar outcomes. In another excellent example [40], the authors were able to detect and quantify fewer than 100 molecules/μL simply by requiring that three antibodies bind to the target simultaneously in order to generate a signal. This equilibrium-based approach greatly reduced the probability of background signal generation and conferred ultra-high resolution. Other digital approaches based on molecular confinement [16], [41] have also illustrated the concepts discussed here, achieving successful quantitation at target concentrations far below the affinity of the binder.

## The Challenge of Specificity

If the detection of analytes at concentrations orders of magnitude below the *K*_D_ is possible, is the effort [42], [43] to develop high affinity reagents unnecessary? This question seems to be particularly salient given the potential performance that can be achieved with a moderate-affinity binder and a focus on decreasing the background signal. However, there remains another important driving force providing the need for high-affinity detection reagents: **specificity**. Every biomolecule has an effective binding affinity to every other biomolecule, even though in most cases this affinity will be exceedingly low. Since most biomolecules can form multiple hydrogen bonds with one another, many off-target interactions will still produce a measurable *K*_D_ for a given affinity reagent, even if this affinity is many orders of magnitude weaker than that for the intended target. Since the concentrations of target molecules can be many orders of magnitude lower than the concentrations of off-target molecules [44], achieving high specificity through biomolecule-mediated molecular recognition is critical. Even if one can still detect low target concentrations with modest affinity binders, the ability to distinguish between target and off-target molecules decreases as the affinity for on-target molecule decreases or as the concentration of off-target molecules increases. This becomes problematic if we do not understand the limitations imposed by imperfect specificity.

Therefore, we would like to conclude this perspective with a brief discussion on the limitations imposed by specificity. Imagine a scenario where an affinity reagent has a *K*_D1_ of 1 μM for a desired target, *T*_1_. Assuming [*T*_1_] ≫ [*A*], the log-linear signal is given by

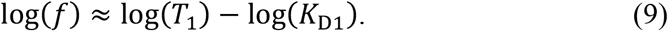

Upon the addition of an off-target molecule, *T*_A_, for which the affinity reagent exhibits an affinity of *K*_D2_, then the total signal is now approximately

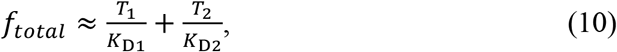

assuming [*T*_2_] ≫ [*A*], [*T*_1_] ≪ *K*_D1_, and [*T*_2_] ≪ *K*_D2_. The log-linear range is now

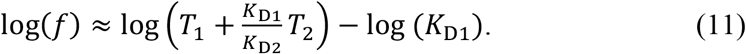

In this scenario, measurements of *T*_1_ in the presence of *T*_2_ will be off by a factor of 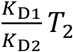 This implies that the impact of off-target specificity becomes problematic if *T*_1_/*T*_2_ does not greatly exceed *K*_D1_/*K*_D2_. This reinforces the need for high affinity reagents. For example, consider the detection of a target that is present at 1 nM in whole blood. Albumin is the off-target protein with the highest concentration in blood (~5 mM) [45]. If albumin binds weakly to the affinity reagent with *K*_D2_ ~500 mM, then *K*_D1_ would need to be less than 9 nM for the error resulting from off-target binding to albumin to be smaller than 10%. As such, the concentrations and cross-reactivity of off-target molecules impose a lower limit on the affinity that is acceptable for an affinity reagent in a given context. This complication is often overlooked, perhaps because addressing it would require the impractical task of profiling each reagent’s *K*_D_ for all possible off-target molecules. Perhaps confirmation bias is also at play; we have a tendency to seek confirming rather than disconfirming information [46]. This bias is exacerbated when unwarranted assumptions—such as the assumption that affinity reagents are highly specific—make disconfirming evidence seem unlikely. Fortunately, specificity can be increased through assay design. For example, by requiring multiple binding events to produce a signal [6], [40], [47], the probability of two binding events occurring on the same off-target molecule is greatly reduced. Therefore, specificity can be improved by requiring dual (or higher [40]) recognition for signal generation, an approach that has the added benefit of reducing background signal and therefore extending the log-linear dynamic range.

## Concluding Remarks

A rigorous understanding of the core principles underlying binding assays is needed to guide the generation of affinity reagents that are best suited for the development of effective biosensors. Some of the conventional wisdom related to binding assays may be stifling the innovation that could be possible with novel technologies. For instance, we have discussed the limiting effects of over-reliance on *K*_D_ as a measure of biosensor performance and how our own biases cause us to overlook important factors like specificity because they are challenging to quantify. By considering binding assays in terms of concentration-dependent shifts in equilibrium, we have embarked on a critical assessment of the key features of these systems, with the goal of identifying important considerations for future biosensor development. First, we explored how the *K*_D_ of a given reagent for its target should only be judged as “better” or “worse” depending on the context in which it is to be employed. We showed that high-resolution measurements can be made even with a low affinity binder, and that by making use of a digital detector (in which binding events are counted rather than averaged in an ensemble) and a low-background assay design, one can extend the log-linear detection range to far lower target concentrations that are independent of *K*_D_ and limited only by Poisson noise. This is in stark contrast to the conventional view that an affinity reagent can only achieve quantitative detection at concentrations with a lower limit of ~*K*_D_/9. This implies that less effort should be spent developing high affinity binders and more on assay development. However, we also show that the upper limit of useful *K*_D_’ for an affinity reagent is imposed by its specificity; higher concentrations of off-target molecules and stronger binding to these molecules necessitates the use of detection reagents with higher affinity for the target.

We conclude by noting that metrics such as *K*_D_, LOD, detection range, and specificity aren’t the end of the story (see Outstanding Questions). Even with perfect molecular quantification, diagnostic assays are still limited by biological variability, which impacts clinical sensitivity and specificity [48]. Although the extent to which assay development should focus on LOD, sensitivity, specificity, or resolution will depend on the context in which the assay is to be used, the design of an optimal assay will inevitably entail achieving the right *K*_D_, a method of background suppression, a digital readout, and dual recognition (or some other mechanism to ensure specificity). It is our hope that the concepts presented here will instill an appreciation of the unexplored opportunities for pushing the diagnostic capabilities of our ever-increasing technological armamentarium.

## GLOSSARY

**Affinity Reagent:** A molecule that binds to a specific target molecule of interest for the purpose of detection or quantification. An affinity reagent can be an antibody, peptide, nucleic acid, or small molecule.

**Aptamer:** A specific type of affinity reagent that is made of DNA or RNA. They are produced through repeated rounds of directed evolution.

**Binding Assay:** A method of molecular quantification that involves the binding of a one molecule to another, i.e. receptor to ligand, antibody to antigen, etc. The addition of target molecule shifts the equilibrium towards a higher fraction of target-bound molecules.

**Binding Curve:** A response function that characterizes the amount of binding as a function of target concentration.

**Detection Range:** The range between the lowest to highest concentrations over which the assay output signal has a large derivative with respect to target concentration. This is often considered to be the concentrations over which the binding curve corresponds to between 10% to 90% of that total signal.

**Dissociation Constant (K_D_):** An equilibrium constant that characterizes the strength of binding between two molecules. It has units of concentration and is inversely proportional to binding affinity.

**Immunoassay:** A method of molecular quantification that utilizes antibody-antigen interactions to achieve detection.

**Limit of Detection (LOD):** The lowest concentration that is statistically discernable from background signal.

**Sensitivity:** In this context, sensitivity refers to the ability of an assay to distinguish between small changes in concentration.

**Specificity:** In this context, specificity refers to the ability of an assay to discriminate between non-cognate and cognate molecules.

## Acknowledgements

We thank Dr. Evelin Sullivan of the Technical Communication Program at Stanford and Mr. Michael Eisenstein for their thoughtful comments and edits on the manuscript.

